# The Evolving Bileome: A Pan-Repository Landscape of Expanded Bile Acid and Related Steroidal Signatures

**DOI:** 10.64898/2026.10.10.758218

**Authors:** Helena Mannochio-Russo, Megan R. Nelson, Esteban Charria-Girón, Laura R. Torres-Ortega, Julius Agongo, Wilhan D. Gonçalves Nunes, Abubaker Patan, Michael Strobel, Lurian C. David, Morgan Panitchpakdi, Emily C. Gentry, Andrés Mauricio Caraballo-Rodríguez, Jasmine Zemlin, Zdenek Kamenik, Nina Acebo, Yessica A. Montoya Giraldo, Anelize Bauermeister, Mingxun Wang, Justin J.J. van der Hooft, Lee R. Hagey, Ipsita Mohanty, Pieter C. Dorrestein

## Abstract

Bile acids and related steroidal metabolites are among the most studied biomolecules, yet recent work has revealed far greater structural diversity than previously recognized, much of it shaped by the microbiome. We previously introduced a repository-scale bile acid MS/MS library built from queries targeting C24 bile acid fragmentation, limited to Orbitrap Q Exactive datasets within GNPS/MassIVE. Here we announce a major expansion of this library: a bile acid and steroidal MS/MS spectral library assembled across three major metabolomics repositories, now capturing C23, C24 (including alkamine), C26 (C–C bonded aceto derivatives), and C27 bile acids, as well as C28 bile alcohols. The expanded libraries comprise 640,141 unique MS/MS spectra, of which 120,293 (18.8%) match 2,529 reference spectra, leaving most of this steroidal chemical space structurally unassigned. Linked to harmonized pan-repository metadata, the library provides a starting point for classifying MS/MS spectra as likely bile acids or related steroids, with two example analyses illustrated here. This resource allows researchers to flag candidate bile acid and steroidal signatures in their own data for further characterization.

## Background & Summary

Bile acids exemplify the paradox of metabolic knowledge: although they were among the first metabolites to be structurally characterized, dating back to 1848^1^, new bile acid analogs and related steroidal metabolites continue to be reported, revealing a broad chemical diversity. These molecules can undergo dozens of structural variations^2^, including hydroxylation, dehydration, epimerization, esterification, sulfation, glucosylation, and, more recently, microbial amidation^3^, alkamination^4^, and C–C-linked aceto-derivative formation^5^. This chemical diversity extends far beyond the canonical C24 bile acids conjugated to glycine and taurine, and as a consequence, a diverse set of tandem mass spectrometry (MS/MS) signatures can be detected in a given MS-based metabolomics analysis.

Despite progress, the systematic annotation of bile acids and related steroids continues to be a major challenge. Many ion forms of existing and yet-to-be-described steroidal molecules are detected in untargeted metabolomics experiments but remain as unannotated features. To help assign the mass spectrometry features that belong to bile acids and related steroidal-derived ions, we previously applied the Mass Spectrometry Query Language (MassQL)^6^ to filter public metabolomics data based on fragmentation patterns characteristic of bile acids. This approach resulted in the creation of a public library of MS/MS spectra that contains diagnostic ions of bile acids and includes different ion forms (ISF and different adducts). This MassQL-generated library comprised of 21,259 MS/MS spectra^7^ that can be used to annotate bile acid MS/MS spectra in metabolomics datasets, and it facilitated the discovery of new bile acids modifications (including amidates of polyamines^7^, drugs^8^, neurotransmitters^9^, and diet-derived molecules^10^) as well as an additional subclass of bile acids belonging to the alkamine class^4^, in which amines react with keto bile acids and steroids to form secondary amines.

These initial repository-scale analyses in GNPS/MassIVE were constrained by computational limitations we faced at that time. The computational infrastructure could not support exhaustive MassQL queries across the full GNPS/MassIVE repository, restricting our analyses to a subset of Orbitrap datasets, specifically those acquired on Q-Exactive instruments. Consequently, Q-TOF MS/MS spectra were not represented in the resulting library. Over the past two years, however, the infrastructure for pan-repository analysis has expanded. Pan-ReDU^11^ now provides indexed, harmonized metadata for more than one million LC-MS/MS files across 5,653 public metabolomics datasets, and advances in indexed repository allow searches using MassQL to be done at larger scales. In addition to these computational constraints, the original library was designed specifically to identify non-to pentahydroxylated C24 bile acids and related steroids. It therefore did not capture several other relevant bile acid-related subclasses, including C23 and C27 bile acids, C24 alkamines, C28 bile alcohols, and C26-derived aceto-variants containing covalent C–C linkages.

Here, we describe a pan-repository MS/MS resource obtained from GNPS/MassIVE^12^, MetaboLights^13^, and Metabolomics Workbench^14^. The resource consists of six spectral libraries representing C23, C24, C24 alkamine, C26, C27, and C28 bile alcohol cores. Diagnostic fragmentation patterns were used to retrieve candidate spectra using MassQL, followed by filtering, spectrum clustering, spectral quality assessment, confidence-aware molecular networking, molecular-formula annotation, and annotation of potential alternative ion forms. Repository provenance was retained, and spectra were linked where possible to harmonized ReDU metadata to provide biological and sample context^11,15^. The resulting libraries are intended as a reusable resource for recognizing candidate bile acid and related steroidal MS/MS signatures and for prioritizing spectra for subsequent structural characterization.

## Methods

### Input data

The spectral libraries were constructed from publicly available LC-MS/MS datasets accessible for repository-scale querying through the GNPS2 infrastructure. Searches were performed across three major metabolomics repositories: GNPS/MassIVE^12^, MetaboLights^13^, and Metabolomics Workbench^14^. The searches were conducted from July/2025 to December/2025. MassQL result tables were retained throughout processing to preserve the relationship between retrieved MS/MS scans and their originating repository datasets, files, and scan numbers. These provenance records were subsequently used to recover precursor information, retention times, and standardized sample metadata using ReDU^15^. The complete ReDU metadata table was downloaded on August 13th, 2025, from https://redu.gnps2.org/.

### MassQL query design

Bile acids and related steroids contain a fused tetracyclic scaffold that yields characteristic fragment ions in collision-induced dissociation. Diagnostic ions for the bile acid classes included in this work (C23, C24, C24 with alkamine substitution, C24 aceto and C28 bile alcohols) were determined manually by inspecting reference MS/MS spectra in existing spectral libraries. The MS/MS spectra for only the 3-aceto dihydroxy C26 bile acid were not available in the GNPS spectral libraries. Therefore, we used the MS/MS spectra available in the publication that discovered the 3-aceto bile acid core for the very first time^5^. The MS/MS spectra in that publication were reported in negative ionization mode, so we converted the diagnostic peaks to the corresponding positive ionization mode by adding the mass of two protons. These diagnostic ions were used to design queries with the Mass Spec Query Language (MassQL)^6^, specifically for positive ionization mode, which are available in **Supplementary Table 1**. These queries were first run against the libraries available in GNPS2 and were refined when necessary to achieve low false discovery rates (details in the **Technical Validation** section). Representative diagnostic fragmentation behavior is shown in **Supplementary Figure 1**.

### Repository-scale MassQL searching

The designed queries were run on GNPS2 against the GNPS/MassIVE, MetaboLights, and Metabolomics Workbench. Because GNPS/MassIVE contains the largest number of MS/MS spectra^11^ compared to the other repositories, querying this entire repository in a single search using the GNPS2 platform exceeded the available computational capacity and prevented the queries from completing. To overcome this limitation, MassQL searches in GNPS/MassIVE were partitioned into three instrument-based collections available in GNPS2: the QE collection, Orbitrap collection, and Q-ToF collection. In MetaboLights and Metabolomics Workbench, the full repositories could be searched with a single GNPS2 job. Therefore, for each bile acid core query, five MassQL jobs were run in GNPS2. A total of 12,388,188 MS/MS spectra were retrieved from all the queries and the repositories combined. Compared to the previous C24 bile acid library^7^, the greater diversity of datasets, instrument platforms, and acquisition conditions represented in these expanded searches motivated us to design a multi-stage filtering and stricter quality-control workflow aiming to select the spectra that most probably represent naturally occurring bile acids.

### Initial filtering of candidate spectra

Retrieved scans were subjected to a series of filtering steps. First, potential duplicate spectra occurring in overlapping GNPS/MassIVE, Q Exactive, and Orbitrap collections were removed. Second, spectra originating from instruments or file structures in which multiple microscans were assigned the same scan number were excluded because these records could lead to ambiguous retrieval of the corresponding MS/MS spectrum. Third, scans without an assigned precursor ion were removed. For datasets where ReDU metadata was available, we used it to remove spectra acquired in data-independent mode, as well as spectra acquired in negative ionization mode. Spectra originating from datasets associated with organic synthetic standards were also removed, as the focus was to retrieve spectra observed in biological samples.

For each retrieved spectrum, a delta mass was defined relative to its corresponding unmodified bile acid or steroidal core. Candidate modifications were retained only when the corresponding delta mass was observed in at least two independent datasets. This filter was not applied to C28 bile alcohols because their queries identified the number of hydroxyl groups on the side chain but did not establish the hydroxylation state of the steroidal core, preventing calculation of a corresponding delta mass. Finally, scans were excluded when the precursor *m/z* was below the highest diagnostic product ion required for the corresponding core query.

### Spectral clustering

Redundancy among the retrieved MS/MS spectra was reduced using the Falcon clustering algorithm^16^ (version 1.3). Clustering was performed with a precursor ion tolerance of 20 ppm, a fragment ion tolerance of 0.02 Da, a maximum cosine distance (eps) of 0.1 (i.e., a minimum cosine similarity of 0.9 between clustered spectra), a minimum of 4 fragment peaks per spectrum, a lower *m/z* bound of 50, and a requirement that spectra be present in at least two samples. For each resulting cluster, the most representative spectrum was retained using Falcon’s --export_representatives option. Specifically, it retains the medoid spectrum, defined as the spectrum with the smallest overall spectral distance to the other spectra in the cluster. This spectrum was used for downstream analyses.

### Spectral entropy calculation

To measure the MS/MS information content, the spectral entropy was calculated for each MS/MS spectrum using classical unweighted Shannon entropy methodology as previously described^17^. For each spectrum, fragment peaks with intensities below 1% of the base peak were removed to reduce spectral noise, retaining all remaining spectral peaks. The intensities of the retained peaks were normalized to a probability distribution,

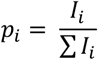

and spectral entropy was calculated as

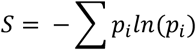

Spectral entropy and fragment-ion count were used as additional measures of spectral quality. Across the bile acid core groups, fragment ion counts increased exponentially at entropy values above approximately 7.5, while MS/MS spectra that matched to bile acids in spectral libraries were largely observed at values below 7.0. We therefore excluded spectra with entropy values >7.5.

### Confidence-aware molecular networking and network-based filtering

The clustered spectra (which refers to the most representative one exported post Falcon clustering - more details on Spectral Clustering section) were further evaluated and filtered using confidence-aware molecular networks generated with SpecReBoot^18^. First, clustered MGF files were preprocessed using DEFAULT_FILTERS from the *matchms* framework^19^ to harmonize spectral metadata. For each bootstrap replicate, fragment ion bins were resampled from the global fragment space, and spectral similarities were recalculated using the modified cosine similarity implemented in FlashSimilarity with a fragment mass tolerance of 0.02 Da. A total of 100 bootstrap replicates were computed for each dataset.

To account for differences in dataset size and network density, the mutual *k*-nearest neighbor parameter (*k*) was adjusted according to the number of spectra in each dataset. Smaller datasets (<2,000 spectra) were processed using *k* = 5-10, intermediate datasets using *k* = 10-20, and larger datasets (>30,000 spectra) using *k* = 30. Dual threshold-based molecular networks were constructed using a minimum mean spectral similarity threshold of 0.7 and a minimum bootstrap edge-support threshold of 0.5.

The clustered spectra from falcon were searched against the GNPS2 public spectral libraries using cosine-based spectral library matching (minimum cosine similarity of 0.7, minimum of 6 matched peaks, precursor and fragment ion tolerances of 0.02 Da), and each match was manually inspected and classified as a bile acid or related steroid (true) or as a compound from another chemical class (false). After the generation of confidence-aware molecular networks, molecular families in which more than one-third of the annotated nodes matched to compounds that are not bile acids were removed in their entirety. In molecular families where one-third or fewer of the annotated nodes were classified as not bile acids, only those individual nodes were removed, while singletons were filtered solely based on their annotation (non-bile acids were removed).

### In silico molecular formula annotation

The molecular formulas of the clustered spectra were generated through two tools: SIRIUS (version 6.2.2)^20^ and BUDDY^21^. For BUDDY, we used the *msbuddy* workflow available in GNPS2 with precursor and fragment ion tolerances of 10 and 20 ppm, respectively, and with halogens enabled. Only molecular formulas predicted with BUDDY with an estimated FDR below 0.05 were considered. For SIRIUS, precursor and fragment ion tolerances were likewise set to 10 and 20 ppm, with the QToF profile specified and [M+H]^+^ as the ion form. Element counts were defined as H[19-], C[17-], N, O[1-], P, and the presence of Cl, Br, and S was set to auto-detection. Only formulas with a SiriusScoreNormalized above 0.9 and at least five fragments explained by SIRIUS were retained.

The predicted molecular formulas from the two tools were reported separately in spectrum-level annotation tables. When both tools returned a formula for an entry, both were retained; no consensus formula was selected or additional ranking based on expected bile acid composition was applied. For each retained prediction, a corresponding delta mass formula was calculated by subtracting the molecular formula of the assigned bile acid core using the *chemparse* package in Python (https://pypi.org/project/chemparse/). These delta mass formulas were added as putative annotations of the modified bile acids in the libraries.

### Estimation of potential alternative ion forms

The generated spectral libraries can contain MS/MS spectra that correspond to different ion forms of the same compound, such as in-source fragments, adducts, and isotopologues. To estimate the proportion of such entries, we reasoned that a single compound entering the ion source can give rise to multiple features, and that two features representing different ion forms of the same compound must therefore (i) differ in precursor *m/z* by a defined exact-mass offset and (ii) co-elute within the same sample.

For every cluster that survived all filtering steps and was retained in the final libraries, we traced back all MS/MS scans that contributed to that cluster and retrieved, from the corresponding MassQL result tables, the originating file path, scan number, precursor *m/z*, and retention time of each scan. Within each file of origin, we compared the retrieved precursors: any two precursors whose *m/z* difference matched one of a set of predefined exact-mass offsets to within 20 ppm and whose retention times agreed within a tolerance of 10 s were flagged as a co-eluting ion-form pair. The offsets considered were ^13^C isotopologues (M+1; +1.003355 Da for singly charged and +0.501678 Da for doubly charged species), sodium (Na-H, +21.981944 Da), potassium (K-H, +37.955882 Da), calcium (Ca-2H, +37.946941 Da), and ammonium (NH_4_-H, +17.026549 Da) adducts, and in-source neutral losses of one to four water molecules (n × 18.010565 Da), carbon monoxide (+27.994915 Da), and carbon dioxide (+43.989830 Da). For isotopologue and adduct offsets, the higher-*m/z* member of the pair was assigned as the alternative ion form and the lower-*m/z* member as the originating species; for in-source neutral losses, the lower-*m/z* member was assigned as the in-source fragment and the higher-*m/z* member as the intact species. A cluster was flagged as a potential alternative ion form if any of its contributing scans participated in such a pair in any file.

These flags were added to the final library annotations, and the corresponding entries were retained rather than removed, as annotations of alternative ion forms are themselves informative. Of the 640,141 entries across all final libraries combined, 88,405 (13.8 %) were flagged as potential alternative ion forms, distributed across isotopologues, adducts, and in-source fragments. The distribution of flagged entries across different ion forms was visualized as Sankey diagrams for each library and for each hydroxylation state within each library. These diagrams were generated with the Plotly package (version 6.3.1) in Python (version 3.10.18).

In addition, the MS/MS spectra in the libraries that are relative to negative values of delta masses were flagged as potential in-source fragmentation. Considering this additional flag, the number of other ion forms increases to 252,470 (39.4%).

### Mapping spectra to harmonized ReDU metadata

Repository provenance was used to associate retained spectra with standardized ReDU sample metadata^11^. ReDU metadata used for this analysis were accessed on August 13, 2025. For each retained spectrum, originating repository file paths were traced through the pre-Falcon annotation tables and matched to ReDU records by dataset accession and file name. Human-related datasets were identified by the presence of “9606|Homo sapiens” in the NCBITaxonomy column, and rodent datasets by the taxonomic identifiers “10088|Mus”, “10090|Mus musculus”, “10105|Mus minutoides”, “10114|Rattus”, or “10116|Rattus norvegicus”. The body part origin of each sample was retrieved from the UBERONBodyPartName field, while entries without an assigned body part were excluded, and only features with a positive delta mass relative to the corresponding bile acid core were retained. For each body part, we then determined, per bile acid core group, the number of contributing MS/MS scans and the number of modifications, defined as unique delta masses (rounded to two decimal places), observed for that group. The results were visualized as bubble plots, generated with Matplotlib (version 3.10.3) in Python (version 3.10.18), separately for human and rodent datasets.

### Taxonomic distribution

The number of retained MS/MS spectra and unique delta masses was also summarized according to the NCBITaxonomy field. Taxonomic information was retrieved from the NCBI Taxonomy database using a custom API key and the taxize package in R (version 4.5.1). Organisms with complete taxonomic lineages (domain, kingdom, phylum, class, order, family, genus, and species) were retained to generate taxonomic trees using the ape and taxize packages. A species-level tree containing organisms across all kingdoms was generated, and additional trees were generated for organisms within the Metazoa kingdom. The number of spectra assigned to individual bile acid subclasses and the total delta masses for each organism were displayed on the taxonomic trees in R (version 4.5.1) using the ape, readr, dplyr, tidyr, purrr, RColorBrewer, and fields packages.

### Use case: vertebrate bile sample collection

The six libraries were evaluated against an independently processed collection of vertebrate gallbladder bile samples available in GNPS/MassIVE under accession MSV000085120. The converted .mzXML files were processed using MZmine 4^22^ (version 4.10.6) and subjected to the feature-based molecular networking workflow^23^ in GNPS2. Library searching was performed against the six libraries generated in this work with a 0.7 cosine similarity threshold, minimum of 6 matched peaks, and precursor and fragment tolerances set to 0.02 Da. Library matches were subsequently filtered with the post-molecular-networking MassQL MetaboApp^24^, retaining only features whose spectra satisfied the list of 25 diagnostic-ion queries designed in this work (**Supplementary Table 1**).

Metadata were restricted to files classified as animal biofluids, all of which represented bile samples in this use case, yielding 1,616 files. NCBI-derived taxonomic assignments were obtained from the accompanying metadata. Because NCBI assigns no class rank to Testudines, Crocodylia, or Lepidosauria, these groups were analyzed as separate lineages. Twenty-one files without a usable class assignment were excluded. The single Coelacanthiformes file, assigned to Actinistia, was also excluded because lineages represented by fewer than three files were not considered. The final class-level analysis comprised 1,594 files from 1,287 species across nine vertebrate lineages.

For each file, peak areas were summed across all validated metabolite features assigned to each bile acid core and divided by the summed peak area of all of the MassQL-validated features, yielding the percentage of the annotated bile acid signal attributed to each core. All ion forms were considered. Specifically, for sample *s* and bile acid core *c*, the relative contribution was calculated as:

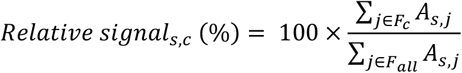

Where *A*_*s,j*_ is the peak area of feature *j* in sample *s, F*_*c*_ is the set of validated features assigned to core *c*, and *F*_*all*_ is the set of all 925 validated features. Thus, the relative contributions of all bile acid cores sum to 100% within each sample. Each scatter represents one LC-MS/MS file, and boxplots show the median and interquartile range, with whiskers extending to 1.5 times the interquartile range. These percentages describe composition within the fraction of the bile salt pool recoverable by the present libraries and queries and do not represent absolute molar composition. Analyses were performed in Python (3.10.18) using pandas (2.3.0), NumPy (2.2.6), and matplotlib (3.10.3).

### Use case: mammalian fecal metabolomics dataset of zoo animals

As a second use case, we applied the spectral libraries to a previously published untargeted LC-MS/MS dataset of fecal samples from zoo mammals, available in GNPS/MassIVE under accession MSV000086131^25^. The converted .mzML files were processed using MZmine 4^22^ (version 4.10.6) and subjected to the feature-based molecular networking workflow^23^ in GNPS2. Library searching was performed against the six libraries generated in this work with 0.7 cosine similarity threshold, minimum of 6 matched peaks, and precursor and fragment tolerances set to 0.02 Da. Library matches were subsequently filtered with the post-molecular-networking MassQL MetaboApp^24^, retaining only features whose spectra satisfied the list of 25 diagnostic-ion queries designed in this work (**Supplementary Table 1**).

For each sample, the relative contribution of each bile acid core to the annotated bile acid signal was calculated as described above for the vertebrate bile dataset. Pairwise comparisons among dietary groups were performed separately for each detected core using two-sided Mann-Whitney U tests. P values were adjusted across all 15 comparisons using the Benjamini-Hochberg procedure. Boxplots show the median and interquartile range, with whiskers extending to 1.5 times the interquartile range; individual scatters represent LC-MS/MS files. Analyses were performed in Python (3.10.18) using pandas (2.3.0), NumPy (2.2.6), SciPy (1.15.3), and matplotlib (3.10.3).

#### Data records

The primary MS/MS resources generated in this study comprise six positive ion mode MS/MS spectral libraries corresponding to C23 bile acids, C24 bile acids, C24 alkamine bile acids, C26 C–C-linked aceto derivatives, C27 bile acids, and C28 bile alcohols. The libraries are provided in.mgf file format and have been integrated into the public GNPS2 MassQL spectral library collection under the following names:

- GNPS-C23-BILE-ACIDS-MODIFICATIONS (4,442 MS/MS spectra)
- GNPS-C24-BILE-ACIDS-MODIFICATIONS (198,128 MS/MS spectra)
- GNPS-C24-ALKAMINE-BILE-ACIDS-MODIFICATIONS (257,490 MS/MS spectra)
- GNPS-C26-BILE-ACIDS-MODIFICATIONS (4,185 MS/MS spectra)
- GNPS-C27-BILE-ACIDS-MODIFICATIONS (66,753 MS/MS spectra)
- GNPS-C28-BILE-ALCOHOL-MODIFICATIONS (109,143 MS/MS spectra)

The libraries are also deposited in Zenodo (doi.org/10.5281/zenodo.21941572). The deposited data include all the GNPS job links used in this work, the final spectral libraries, and MassQL query definitions, spectrum-level annotation tables, provenance information linking library entries to their source repository files and scans, predicted molecular formulas and delta mass annotations, potential alternative ion form assignments, and harmonized ReDU metadata where available.

The original LC-MS/MS files from which the spectral libraries were constructed remain available through their respective source repositories (GNPS/MassIVE, MetaboLights, and Metabolomics Workbench). Repository provenance retained in the deposited annotation tables allows individual library entries to be traced to their originating dataset, file, and MS/MS scan through a Universal Spectrum Identifier (USI).

#### Data overview

The six final libraries comprise 640,141 MS/MS spectra spanning six classes: C23 bile acids, C24 bile acids, C24 alkamine bile acids, C26 bile acids, C27 bile acids, and C28 bile alcohols (**Figure 1a**). The C24 resource alone expanded from 21,259 spectra in the previously reported library^7^ to 198,128 spectra. Across all six libraries, 120,293 spectra (18.8%) matched 2,529 reference spectra of known compounds using the standard GNPS2 library-search workflow; inclusion of an additional spectral library generated by multiplexed synthesis^26^ increased the number of matched spectra to 135,610 (21.2%). For the C24 class, 8,993 unique delta masses were observed, of which 6,310 remained after excluding entries attributed to potential isotopologues, adducts, or in-source fragments. ReDU-linked metadata contextualize the retained spectra across human and rodent body parts (**Figure 1b, Supplementary Figures 2 and 3**) and across taxa represented in public repositories (**Figure 1c, Supplementary Figure 4**). Applying these libraries to the investigation of a dataset of bile samples from vertebrates showed that C24 signatures comprised the largest relative contribution in most represented vertebrate lineages, whereas Crocodylia, Testudines, and Amphibia showed a larger relative contribution of C27 signatures (**Figure 2a**). In the zoo-mammal fecal dataset, the application of these libraries revealed greater relative contributions of C23, C27, and C28 bile alcohol signatures in herbivores and a higher relative contribution of C24 alkamine signatures in the omnivore and carnivore diet groups (**Figure 2b**). Notably, the subclasses added in this expansion showed diet-associated patterns distinct from, and in several cases more pronounced than, those of canonical C24 bile acids, which would not have been captured by the previous C24-only library. These differences may reflect host physiology, microbial metabolism, or dietary steroid inputs (e.g., plant-derived sterols). These application summaries are descriptive and should not be interpreted as absolute bile acid abundance or as causal effects of host taxonomy or diet.

**Figure 1.**
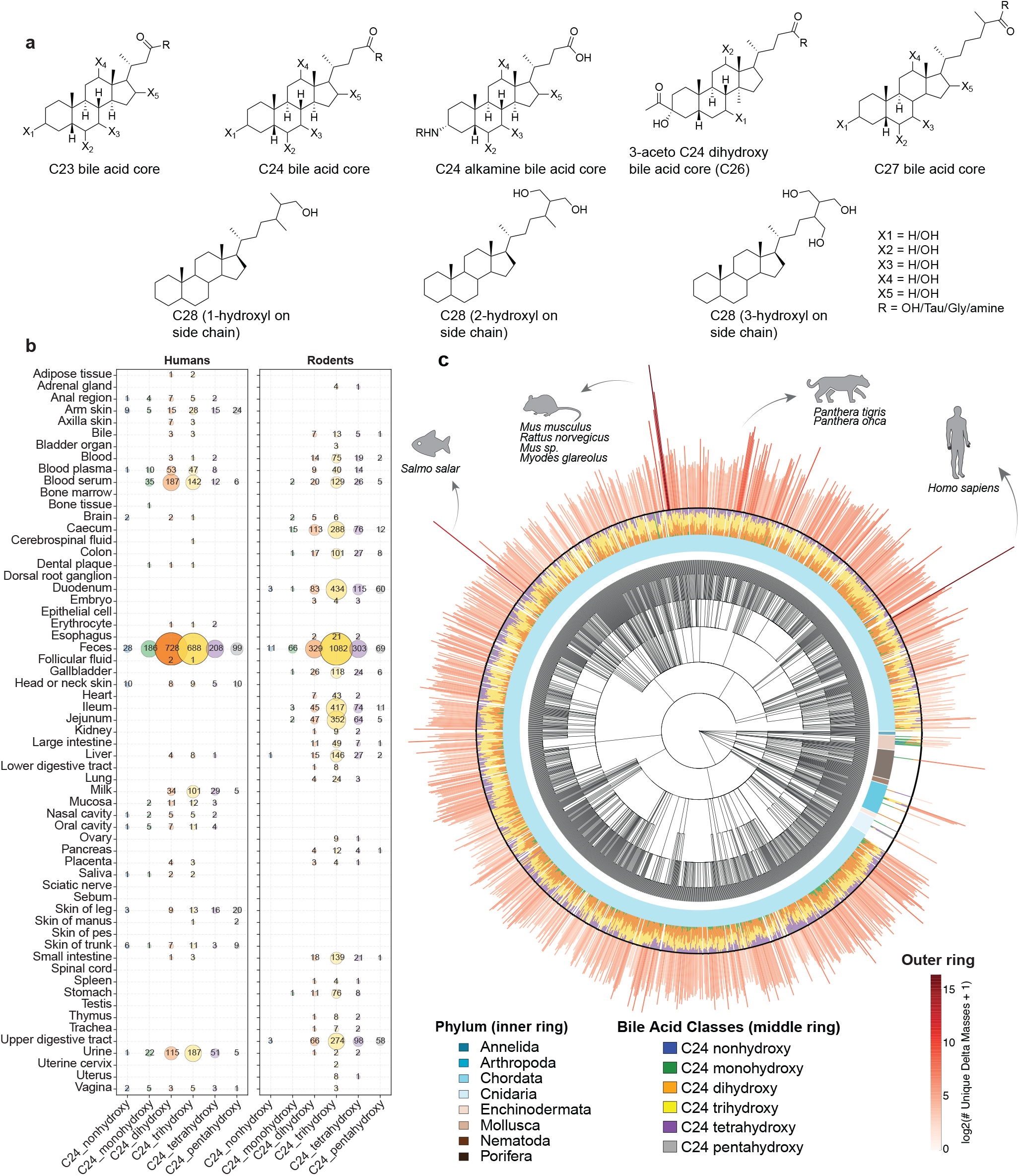
Expanded bile acid spectral libraries map steroidal signatures across biological contexts. **(a)** Representative structures of the bile acid and related steroidal cores used to design MassQL queries: C23, C24, C24 alkamine, 3-aceto C24 dihydroxy (C26), C27, and C28 bile alcohols containing one to three side-chain hydroxyl groups. X_1_-X_5_ indicate variable H/OH substitutions, and R indicates OH, taurine, glycine, amines, or other substitutions. **(b)** Distribution of non-to pentahydroxylated C24 signatures across human and rodent tissues and biofluids identified using harmonized ReDU metadata^11,15^. Values indicate the number of unique delta masses, interpreted as potential modifications, for each C24 bile acid core. Only positive deltas were included, and spectra not assigned as potential in-source fragments, adducts, or isotopes were considered. **(c)** Taxonomic distribution of C24 signatures across organisms within the Metazoa kingdom represented in the repositories. The central tree shows NCBI taxonomic relationships; the inner colored ring denotes phylum, and stacked barplots indicate detected C24 hydroxylation classes. The outer plot represents the number of unique delta masses observed for each species on a log2 scale. Selected animals are highlighted. Apparent differences in bile acid diversity among taxa may also reflect unequal sampling depth and data availability. Icons were obtained from Bioicons.com

**Figure 2.**
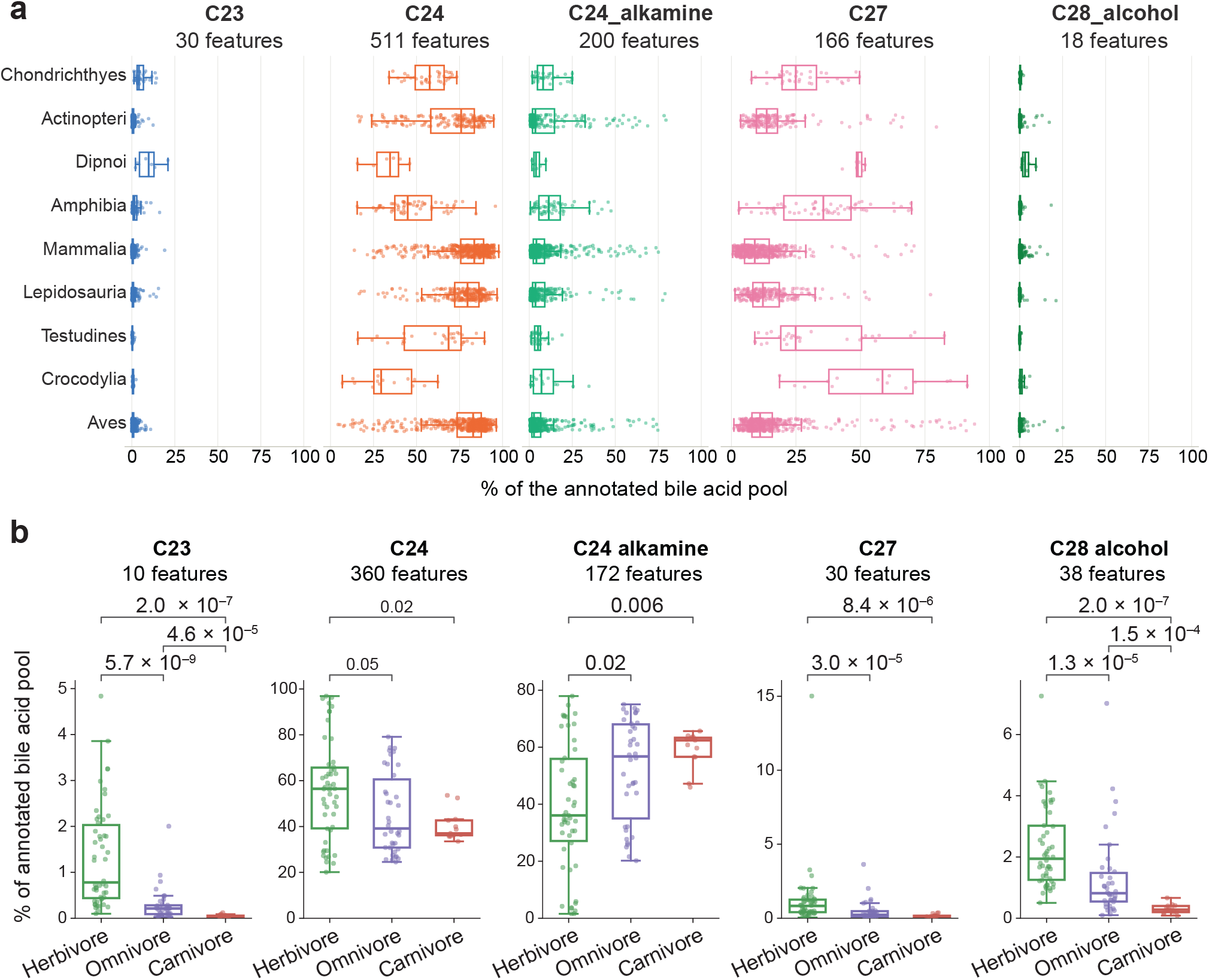
Illustrative applications of the spectral libraries in metabolomics datasets. **(a)** Relative contributions of C23, C24, C24 alkamine, C27, and C28 bile alcohol signatures to the annotated bile acid signal in 1,594 vertebrate bile files across nine lineages (MSV000085120). **(b)** Relative core composition in 101 mammalian fecal samples categorized as herbivores (n = 50), omnivores (n = 38), or carnivores (n = 13). Values above brackets are Benjamini-Hochberg-adjusted p-values from two-sided Mann-Whitney U tests. Center lines show medians, boxes show interquartile ranges, whiskers extend to 1.5 times the interquartile range, and points represent individual files. Percentages describe relative composition of the annotated MS/MS features and not absolute molar abundance.

#### Technical validation

##### Specificity of the queries

The specificity of the MassQL queries was assessed before repository-scale deployment by searching them against the GNPS public spectral libraries, comprising 985,492 MS/MS spectra from diverse compound classes. Each retrieved spectrum was classified as a true or false match based on the structure annotation of the corresponding library entry. Spectra annotated as compounds containing a steroidal tetracyclic core were considered true matches, whereas spectra annotated as compounds from other chemical classes were considered false matches. The FDR of each query was estimated as the percentage of false matches among all evaluated matches. Queries with a high proportion of false matches were refined iteratively by adding diagnostic ions, narrowing m/z tolerances, or introducing minimum relative-intensity thresholds, and re-evaluated until the proportion of false matches was reduced. The final queries used for the generation of the bile acid libraries had FDRs ranging from 0 to 0.035%.

##### Spectral quality and redundancy control

Multiple independent procedures were used to limit low-quality or redundant library entries. Spectra without reliable precursor assignments, spectra associated with data-independent or negative-mode acquisition, duplicated MS/MS spectra, and spectra from organic synthesis datasets were removed before clustering. Falcon^16^ clustering subsequently reduced repeated observations of closely related spectra, with a requirement for representation in at least two samples. Spectral entropy^17^ and fragment-ion counts provided additional quality descriptors (**Supplementary Figure 5**). Library entries may also include chimeric MS/MS spectra containing fragments from co-isolated precursors. For example, a MassQL query for polyethylene glycol (PEG) signatures^27^ retrieved 80 spectra that also contained bile acid diagnostic ions in the C23 and C26 libraries (16 and 64 scans, respectively).

##### Confidence-aware molecular network validation

After clustering, confidence-aware molecular networking provided an additional check on whether MS/MS spectra occupied spectral neighborhoods consistent with the intended steroidal classes. To achieve this, the spectra were searched against the GNPS2 spectral libraries, and each resulting match was manually inspected. Matches to reference compounds containing a tetracyclic steroidal core were treated as supporting annotations, whereas matches to unrelated compound classes were considered evidence of potential false-positive retrieval. The FDR at this stage was estimated, for each library, as the percentage of annotated spectra matching non-steroidal compounds, and ranged from 0.02 to 2.38%. Network-level and node-level pruning was then applied according to the proportion of non-target reference annotations, as described in the Methods section (**Supplementary Figure 6**). This step removed nodes and spectral neighbors inconsistent with the targeted bile acid and related steroidal fragmentation space.

##### Comparison with reference spectra

To assess how much of the libraries corresponds to previously characterized compounds, all library entries were searched against the GNPS2 default reference libraries using the GNPS2 library search workflow, and, separately, against a multiplexed-synthesis library^27^ that contains numerous bile acid conjugates. The resulting matches were predominantly to bile acids and steroidal structures, with 96.1% of matched reference spectra annotated as compounds containing a steroidal core. The remaining unmatched spectra do not have direct reference-spectrum annotations but can still support class-level assignment of candidate bile acid analogs and related steroidal molecules.

##### Evaluation of potential alternative ion forms

Co-elution and exact-mass-offset analysis identified 88,405 of the 640,141 final entries (13.8%) as potential alternative ion forms, including isotopologues, adducts, and in-source fragments. The fraction varied among individual libraries and subclasses (**Supplementary Figures 7 and 8**). Because in-source fragments can also appear as precursor ions with negative delta masses relative to the assigned bile acid core, entries with negative delta masses were additionally flagged. Incorporating this broader criterion resulted in 252,470 entries (39.4% of the combined libraries) carrying a potential alternative ion form or in-source fragment flag. These spectra were retained in the libraries and annotated accordingly, allowing users to recognize and account for such features in downstream analyses.

##### Post-molecular networking MassQL validation

The value of applying core-diagnostic ion rules after conventional spectral library searching was illustrated using two independent use cases. In the first, a vertebrate bile dataset was searched against the libraries in GNPS2, yielding 1,704 candidate bile acid matches. These matches were subsequently evaluated using the 25 MassQL queries developed in this study and used to construct the libraries. Of the initial matches, 779 failed to be retrieved through the queries, whereas 925 were retained. This post-molecular-networking MassQL step, implemented through a GNPS2 MetaboApp^24^, therefore removed candidate matches that exceeded the global cosine-similarity threshold but did not reproduce the core-specific fragmentation pattern required by the original queries. This distinction is important because some diagnostic ions are relatively low in abundance and consequently contribute only modestly to the overall cosine similarity score.

The same workflow was applied to a second dataset comprising mammalian fecal samples, resulting in 610 features that passed the post-molecular-networking MassQL criteria. Together, these use cases illustrate how the spectral libraries can be used as an initial screening tool, followed by the corresponding MassQL queries as a second-stage check for the presence of core-diagnostic fragmentation.

#### Usage Notes

Several points should be considered when using these libraries. Although the libraries were curated to retain spectra with bile acid and related steroidal fragmentation patterns, searching experimental data against them can still yield matches that are not supported by these diagnostic fragmentations. Spectral matching is based on cosine similarity across all fragment ions; consequently, a spectrum can exceed the matching threshold even when a diagnostic ion required by the original MassQL query is absent in the experimental spectrum. Some queries also depend on relative fragment-ion intensities or lower-abundance ions that are not explicitly evaluated by cosine similarity. Within GNPS2, this step can be performed using the post-molecular-networking MassQL MetaboApp^24^.

A match to one of these libraries supports assignment of a steroidal core class and the presence of a candidate modification, but it does not independently establish the modification’s chemical identity, attachment site, regiochemistry, or stereochemistry. Confirmation of a specific conjugate or structure requires orthogonal evidence, such as retention time and MS/MS agreement with an authentic standard or isolation followed by structure elucidation. Similarly, molecular formulas predicted with SIRIUS or BUDDY and formulas assigned to delta masses should be interpreted as candidate annotations. They reduce the possible chemical space associated with an observed mass difference but do not establish a unique chemical transformation. Library entries flagged as potential isotopologues, adducts, or in-source fragments were intentionally retained because they can assist in the recognition of alternative ion forms. However, users are encouraged to assess co-elution as well as for chimerism within their own datasets to determine whether a flagged feature is consistent with an alternative ion form of another detected feature.

Finally, the coverage of these libraries is limited by the diagnostic fragments used to design the queries, positive ionization mode, parameters used for data acquisition, and datasets available at the time the searches were performed. Compounds that preferentially ionize in the negative ionization mode or occur in studies that fall outside the searched repository collections may therefore be absent. Future expansions could use unsupervised approaches such as MS2LDA 2.0^28^ to identify recurring fragmentation patterns and guide the selection of additional diagnostic fragments for MassQL queries.

## Data availability

The six bile acid and related steroidal spectral libraries are publicly available through the GNPS MassQL spectral library collection under the names GNPS-C23-BILE-ACIDS-MODIFICATIONS, GNPS-C24-BILE-ACIDS-MODIFICATIONS, GNPS-C24-ALKAMINE-BILE-ACIDS-MODIFICATIONS, GNPS-C26-BILE-ACIDS-MODIFICATIONS, GNPS-C27-BILE-ACIDS-MODIFICATIONS and GNPS-C28-BILE-ALCOHOL-MODIFICATIONS.

These libraries are also found in Zenodo, together with the annotation and provenance tables: doi.org/10.5281/zenodo.21941572. The following results were also deposited in Zenodo: SIRIUS outputs (doi.org/10.5281/zenodo.23006535), SpecReBoot results (doi.org/10.5281/zenodo.23199804), Falcon clustering results (doi.org/10.5281/zenodo.23219482). The datasets used as use cases are available from GNPS/MassIVE under accession numbers MSV000085120 and MSV000086131, and their feature-based molecular networking job links in GNPS2 are https://gnps2.org/status?task=ea3628cec9d042358b558f86edc781c1 and https://gnps2.org/status?task=3d18cb7996ce4e478a52ec2ab1c80f9d, respectively.

MassQL queries are available in **Supplementary Table 1**; their FDR estimations and GNPS2 job links used to run the queries against the GNPS2 libraries, queries against the repositories, BUDDY molecular formula predictions, and library searches used in this work are also deposited in Zenodo.

## Code availability

The code to perform all the MassQL results filtering steps is available at https://github.com/helenamrusso/library_generation_from_MassQL. The scripts used for estimating different ion forms of MS/MS spectra, for body part and organism distribution, and for the re-analyses of public datasets are available at https://github.com/helenamrusso/bile_acid_updated_library.

## Author contributions

H.M-R. and I.M. performed repository-scale data retrieval. H.M-R., M.R.N., W.D.G.N., and I.M. performed quality-control analyses. H.M-R. and W.D.G.N. performed data filtering. H.M-R. performed molecular formula annotation and alternative ion-form analyses. H.M-R. and L.C.D. performed analysis of the use-case examples. E.C-G., L.R.T-O., and J.J.J.vdH. performed confidence-aware molecular networking. M.R.N. performed taxonomic distribution analyses. I.M. and L.R.H. designed the queries. J.A. synthesized C24 alkamine bile acids to support query design. M.P., E.C.G., Y.A.M.G., Z.K., N.A., A.B., and A.M.C.-R. acquired the vertebrate animal dataset. M.S. and M.W. sourced additional metadata and incorporated the libraries into GNPS2. P.C.D. acquired funding and supervised the project. All authors reviewed the manuscript.

## Competing interests

P.C.D. is an advisor and holds equity in Cybele, BileOmix and Sirenas and a Scientific co-founder, advisor and holds equity to Ometa, Enveda, and Arome with prior approval by UC-San Diego. P.C.D. also consulted for DSM animal health in 2023. M.W. is a co-founder of Ometa Labs LLC. J.J.J.vdH. is a member of the Scientific Advisory Board of NAICONS Srl., Milano, Italy and consults for Corteva Agriscience, Indianapolis, IN, USA.

In the Dorrestein Lab, the use of AI, generative AI, and large language models (LLMs), and software that uses these technologies, both free and commercial, is encouraged across all aspects of research, including literature review, coding, data analysis, and text editing. The MassQL design and library creation were done without AI. All figures and analyses are original. All AI-assisted text was reviewed, verified against the underlying data and results, and edited by the authors, who take full responsibility for the content of the manuscript. For transparency and reproducibility, raw data, data tables, and final code used in this research are made accessible and linked with this manuscript.

## Acknowledgements

H.M-R. and P.C.D. are supported by NIDDK (U24DK133658). L.C.D. is supported by the Fulbright U.S. Student Program through the DDRA (Doctoral Dissertation Research Award), which is sponsored by the U.S. Department of State and the Fulbright Brazil Commission. L.R.T.-O. and J.J.J.vdH. gratefully recognize the Marie Skłodowska-Curie grant under the European Union’s Horizon Europe programme MAGiC-MOLFUN (grant no. 101072485). Z.K. was supported by the Programme Johannes Amos Comenius (grant no. CZ.02.01.01/00/22_008/0004597) of the Ministry of Education, Youth and Sports of the Czech Republic.

