## Supplementary Material for "The Evolving Bileome: A Pan-Repository Landscape of Expanded Bile Acid and Related Steroidal Signatures"

**Addresses:**
[1] Skaggs School of Pharmacy and Pharmaceutical Sciences, University of California San Diego, La Jolla, CA, USA
[2] Bioinformatics Group, Wageningen University & Research, Droevendaalsesteeg 1, 6708 PB Wageningen, the Netherlands
[3] Department of Computer Science and Engineering, University of California, Riverside, Riverside, CA 92521, USA
[4] Department of Chemistry, Virginia Tech, Blacksburg, VA 24061, USA
[5] Institute of Microbiology of the Czech Academy of Sciences, Prague, Czechia
[6] Instituto de Química, Universidade de São Paulo, São Paulo 05508-900, Brazil
[7] Department of Biochemistry, University of Johannesburg, Johannesburg 2006, South Africa
[8] Department of Medicine, University of California San Diego, La Jolla, California 92093, United States
[9] Department of Veterinary and Biomedical Sciences, The Pennsylvania State University, University Park, PA, United States
[10] Department of Nutritional Sciences, The Pennsylvania State University, University Park, PA, United States
[11] Center for Microbiome Innovation, University of California, San Diego, La Jolla, CA, 92093, USA
[12] Collaborative Mass Spectrometry Innovation Center, Skaggs School of Pharmacy and Pharmaceutical Sciences, University of California San Diego, La Jolla, CA, USA
[13] Department of Pharmacology, University of California San Diego, La Jolla, CA, 92093, USA
[14] Laboratório de Métodos de Extração e Separação (LAMES), Instituto de Química (IQ), Universidade Federal de Goiás (UFG), Campus II – Samambaia, Goiânia, GO, 74045-155, Brazil
[15] Research Group in Biosciences and Biotechnology (TechLife), School of Applied Sciences and Engineering, EAFIT University, Medellín, Colombia.

**Correspondence:**
Ipsita Mohanty,
Pieter C. Dorrestein,

### Supplementary Tables

#### Supplementary Table 1. MassQL queries used to retrieve bile acid and related steroidal MS/MS spectra, by core class.

| Bile acid core | Query |
|---|---|
| C24 - Nonhydroxy | QUERY scaninfo(MS2DATA) WHERE MS2PROD=343.30:TOLERANCEMZ=0.01:INTENSITYPERCENT=5 AND MS2PROD=325.29:TOLERANCEMZ=0.01:INTENSITYPERCENT=5 |
| C24 - Monohydroxy | QUERY scaninfo(MS2DATA) WHERE MS2PROD=341.28:TOLERANCEMZ=0.01:INTENSITYPERCENT=5 AND MS2PROD=323.27:TOLERANCEMZ=0.01:INTENSITYPERCENT=5 |
| C24 - Dihydroxy | QUERY scaninfo(MS2DATA) WHERE MS2PROD=339.27:TOLERANCEMZ=0.01:INTENSITYPERCENT=5 AND MS2PROD=321.26:TOLERANCEMZ=0.01:INTENSITYPERCENT=5 |
| C24 - Trihydroxy | QUERY scaninfo(MS2DATA) WHERE MS2PROD=337.25:TOLERANCEMZ=0.01:INTENSITYPERCENT=5 AND MS2PROD=319.24:TOLERANCEMZ=0.01:INTENSITYPERCENT=5 |
| C24 - Tetrahydroxy | QUERY scaninfo(MS2DATA) WHERE MS2PROD=335.24:TOLERANCEMZ=0.01:INTENSITYPERCENT=5 AND MS2PROD=317.23:TOLERANCEMZ=0.01:INTENSITYPERCENT=5 |
| C24 - Pentahydroxy | QUERY scaninfo(MS2DATA) WHERE MS2PROD=333.22:TOLERANCEMZ=0.01:INTENSITYPERCENT=5 AND MS2PROD=315.21:TOLERANCEMZ=0.01:INTENSITYPERCENT=5 |
| C23 - Monohydroxy | QUERY scaninfo(MS2DATA) WHERE MS2PROD=327.27:TOLERANCEMZ=0.01:INTENSITYPERCENT=2 AND MS2PROD=309.26:TOLERANCEMZ=0.01:INTENSITYPERCENT=2 AND MS2MZ = 309.26:TOLERANCEPPM=20:INTENSITYMATCH=Y:INTENSITYMATCHREFERENCE AND MS2MZ=285.25:TOLERANCEPPM=20:INTENSITYMATCH=Y*1.9:INTENSITYMATCHPERCENT=40 |
| C23 - Dihydroxy | QUERY scaninfo(MS2DATA) WHERE MS2PROD=325.25:TOLERANCEMZ=0.01:INTENSITYPERCENT=2 AND MS2PROD=307.24:TOLERANCEMZ=0.01:INTENSITYPERCENT=2 AND MS2MZ = 307.24:TOLERANCEPPM=20:INTENSITYMATCH=Y:INTENSITYMATCHREFERENCE AND MS2MZ=283.24:TOLERANCEPPM=20:INTENSITYMATCH=Y*1.9:INTENSITYMATCHPERCENT=40 |
| C23 - Trihydroxy | QUERY scaninfo(MS2DATA) WHERE MS2PROD=323.24:TOLERANCEMZ=0.01:INTENSITYPERCENT=2 AND MS2PROD=305.23:TOLERANCEMZ=0.01:INTENSITYPERCENT=2 AND MS2MZ = 305.23:TOLERANCEPPM=20:INTENSITYMATCH=Y:INTENSITYMATCHREFERENCE AND MS2MZ=281.23:TOLERANCEPPM=20:INTENSITYMATCH=Y*1.9:INTENSITYMATCHPERCENT=40 |
| C23 - Tetrahydroxy | QUERY scaninfo(MS2DATA) WHERE MS2PROD=321.22:TOLERANCEMZ=0.01:INTENSITYPERCENT=2 AND MS2PROD=303.21:TOLERANCEMZ=0.01:INTENSITYPERCENT=2 AND MS2MZ = 303.21:TOLERANCEPPM=20:INTENSITYMATCH=Y:INTENSITYMATCHREFERENCE AND MS2MZ=279.22:TOLERANCEPPM=20:INTENSITYMATCH=Y*1.9:INTENSITYMATCHPERCENT=40 |
| C23 - Pentahydroxy | QUERY scaninfo(MS2DATA) WHERE MS2PROD=319.21:TOLERANCEMZ=0.01:INTENSITYPERCENT=2 AND MS2PROD=301.19:TOLERANCEMZ=0.01:INTENSITYPERCENT=2 AND MS2MZ = 301.19:TOLERANCEPPM=20:INTENSITYMATCH=Y:INTENSITYMATCHREFERENCE AND MS2MZ=277.19:TOLERANCEPPM=20:INTENSITYMATCH=Y*1.9:INTENSITYMATCHPERCENT=40 |
| C27 - Monohydroxy | QUERY scaninfo(MS2DATA) WHERE MS2PROD=383.33:TOLERANCEPPM=20:INTENSITYPERCENT=5 AND MS2PROD=159.12:TOLERANCEPPM=20:INTENSITYPERCENT=2 |
| C27 - Dihydroxy | QUERY scaninfo(MS2DATA) WHERE MS2PROD=381.32:TOLERANCEPPM=20:INTENSITYPERCENT=5 AND MS2PROD=159.12:TOLERANCEPPM=20:INTENSITYPERCENT=2 |
| C27 - Trihydroxy | QUERY scaninfo(MS2DATA) WHERE MS2PROD=379.29:TOLERANCEPPM=20:INTENSITYPERCENT=5 AND MS2PROD=159.12:TOLERANCEPPM=20: INTENSITYPERCENT=2 |
| C27 - Tetrahydroxy | QUERY scaninfo(MS2DATA) WHERE MS2PROD=377.28:TOLERANCEPPM=20:INTENSITYPERCENT=5 AND MS2PROD=159.12:TOLERANCEPPM=20:INTENSITYPERCENT=2 |
| C27 - Pentahydroxy | QUERY scaninfo(MS2DATA) WHERE MS2PROD=375.27:TOLERANCEPPM=20:INTENSITYPERCENT=5 AND MS2PROD=159.12:TOLERANCEPPM=20:INTENSITYPERCENT=2 |
| C26 aceto dihydroxy | QUERY scaninfo(MS2DATA) WHERE MS2PROD=347.294:TOLERANCEPPM=20:INTENSITYPERCENT=2 AND MS2PROD=329.284:TOLERANCEPPM=20:INTENSITYPERCENT=2 |

| | |
|---|---|
| C24 alkylamine - Monohydroxy | QUERY scaninfo(MS2DATA) WHERE MS2PROD=359.2945:TOLERANCEPPM=20:INTENSITYPERCENT=4 AND MS2PROD=341.2839:TOLERANCEPPM=20:INTENSITYPERCENT=2 AND MS2MZ = 341.2839:TOLERANCEPPM=20:INTENSITYMATCH=Y:INTENSITYMATCHREFERENCE AND MS2MZ= 359.2945:TOLERANCEPPM=20:INTENSITYMATCH=Y*15:INTENSITYMATCHPERCENT=88 |
| C24 alkylamine - Dihydroxy | QUERY scaninfo(MS2DATA) WHERE MS2PROD=357.2788:TOLERANCEPPM=20:INTENSITYPERCENT=4 AND MS2PROD=339.2682:TOLERANCEPPM=20:INTENSITYPERCENT=2 AND MS2MZ = 339.2682:TOLERANCEPPM=20:INTENSITYMATCH=Y:INTENSITYMATCHREFERENCE AND MS2MZ= 357.2788:TOLERANCEPPM=20:INTENSITYMATCH=Y*15:INTENSITYMATCHPERCENT=88 |
| C24 alkylamine - Trihydroxy | QUERY scaninfo(MS2DATA) WHERE MS2PROD=355.2632:TOLERANCEPPM=20:INTENSITYPERCENT=4 AND MS2PROD=337.2526:TOLERANCEPPM=20:INTENSITYPERCENT=2 AND MS2MZ = 337.2526:TOLERANCEPPM=20:INTENSITYMATCH=Y:INTENSITYMATCHREFERENCE AND MS2MZ=355.2632:TOLERANCEPPM=20:INTENSITYMATCH=Y*15:INTENSITYMATCHPERCENT=88 |
| C24 alkylamine - Tetrahydroxy | QUERY scaninfo(MS2DATA) WHERE MS2PROD=353.2475:TOLERANCEPPM=20:INTENSITYPERCENT=4 AND MS2PROD=335.2369:TOLERANCEPPM=20:INTENSITYPERCENT=2 AND MS2MZ = 335.2369 :TOLERANCEPPM=20:INTENSITYMATCH=Y:INTENSITYMATCHREFERENCE AND MS2MZ= 353.2475 :TOLERANCEPPM=20:INTENSITYMATCH=Y*15:INTENSITYMATCHPERCENT=88 |
| C24 alkylamine - Pentahydroxy | QUERY scaninfo(MS2DATA) WHERE MS2PROD=351.2319:TOLERANCEPPM=20:INTENSITYPERCENT=4 AND MS2PROD=333.2213:TOLERANCEPPM=20:INTENSITYPERCENT=2 AND MS2MZ = 333.2213:TOLERANCEPPM=20:INTENSITYMATCH=Y:INTENSITYMATCHREFERENCE AND MS2MZ= 351.2319:TOLERANCEPPM=20:INTENSITYMATCH=Y*15:INTENSITYMATCHPERCENT=88 |
| C28 alcohol - 1OH | QUERY scaninfo(MS2DATA) WHERE MS2PROD=125.13:TOLERANCEPPM=20:INTENSITYPERCENT=10 AND MS2PROD=143.14:TOLERANCEPPM=20:INTENSITYPERCENT=10 |
| C28 alcohol - 2OH | QUERY scaninfo(MS2DATA) WHERE MS2PROD=123.12:TOLERANCEPPM=30:INTENSITYPERCENT=10 AND MS2PROD=141.13:TOLERANCEPPM=30:INTENSITYPERCENT=10 |
| C28 alcohol - 3OH | QUERY scaninfo(MS2DATA) WHERE MS2PROD=121.10:TOLERANCEPPM=20:INTENSITYPERCENT=10 AND MS2PROD=139.11:TOLERANCEPPM=20:INTENSITYPERCENT=10 |

#### Supplementary Figures

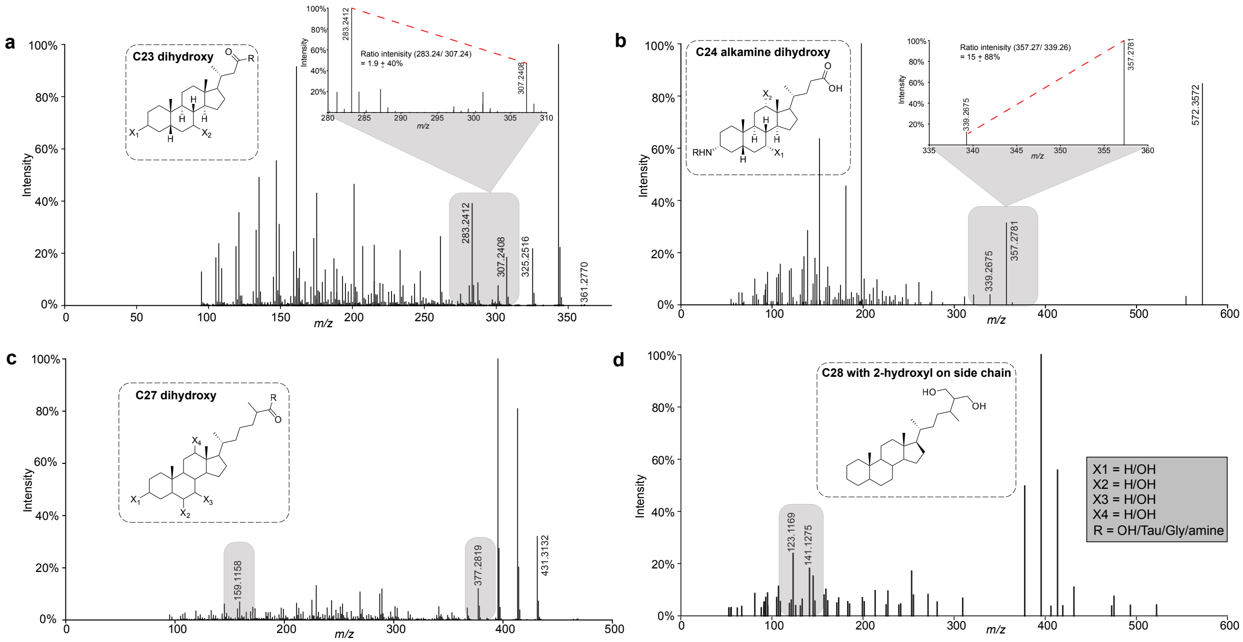


**Supplementary Figure 1. Representative bile acid-inspired MS/MS fragmentation behavior used for MassQL query design.** Positive-ion MS/MS spectra are shown for representative **(a)** C23 dihydroxy bile acid, **(b)** C24 dihydroxy alkamine, **(c)** C27 dihydroxy bile acid, and **(d)** C28 bile alcohol. Gray regions highlight diagnostic fragment ions used to construct and refine MassQL queries. Insets show the corresponding generalized steroidal scaffolds and variable substitution positions.

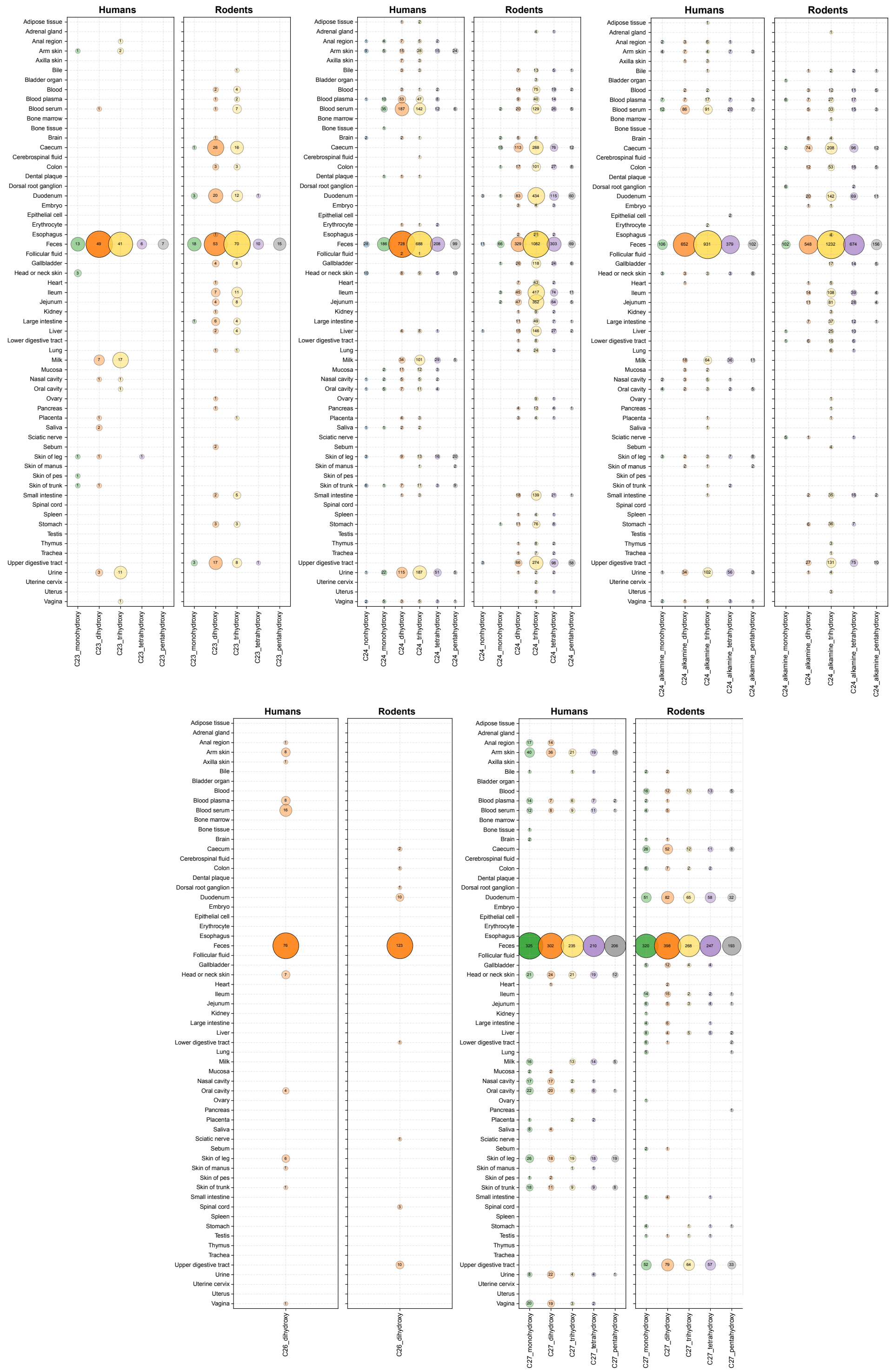


**Supplementary Figure 2. Distribution of bile acid modifications across human and rodent tissues and biofluids.** Bubble plots show the number of unique positive delta masses associated (and not assigned as other ion forms) with each bile acid core and hydroxylation subclass across body parts recorded in the harmonized ReDU metadata. Delta masses were rounded to two decimal places. Values represent the number of unique delta masses; colors distinguish hydroxylation subclasses.

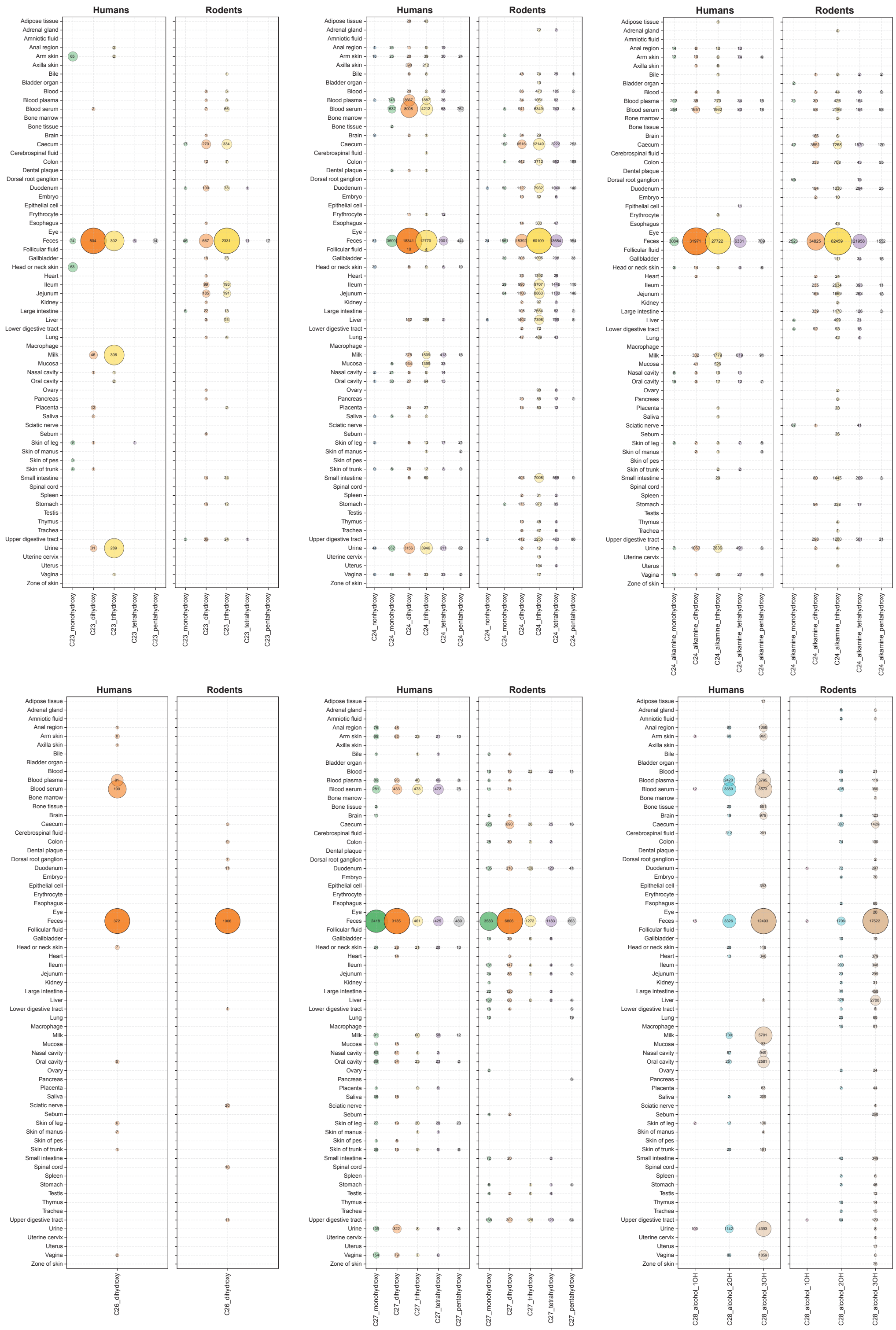


**Supplementary Figure 3. Distribution of bile acid MS/MS scans across human and rodent tissues and biofluids.** Bubble plots show the number of MS/MS scans associated with positive delta masses for each bile acid core and hydroxylation subclass across body parts recorded in the harmonized ReDU metadata. Values within each bubble represent the number of MS/MS scans; colors distinguish hydroxylation subclasses.

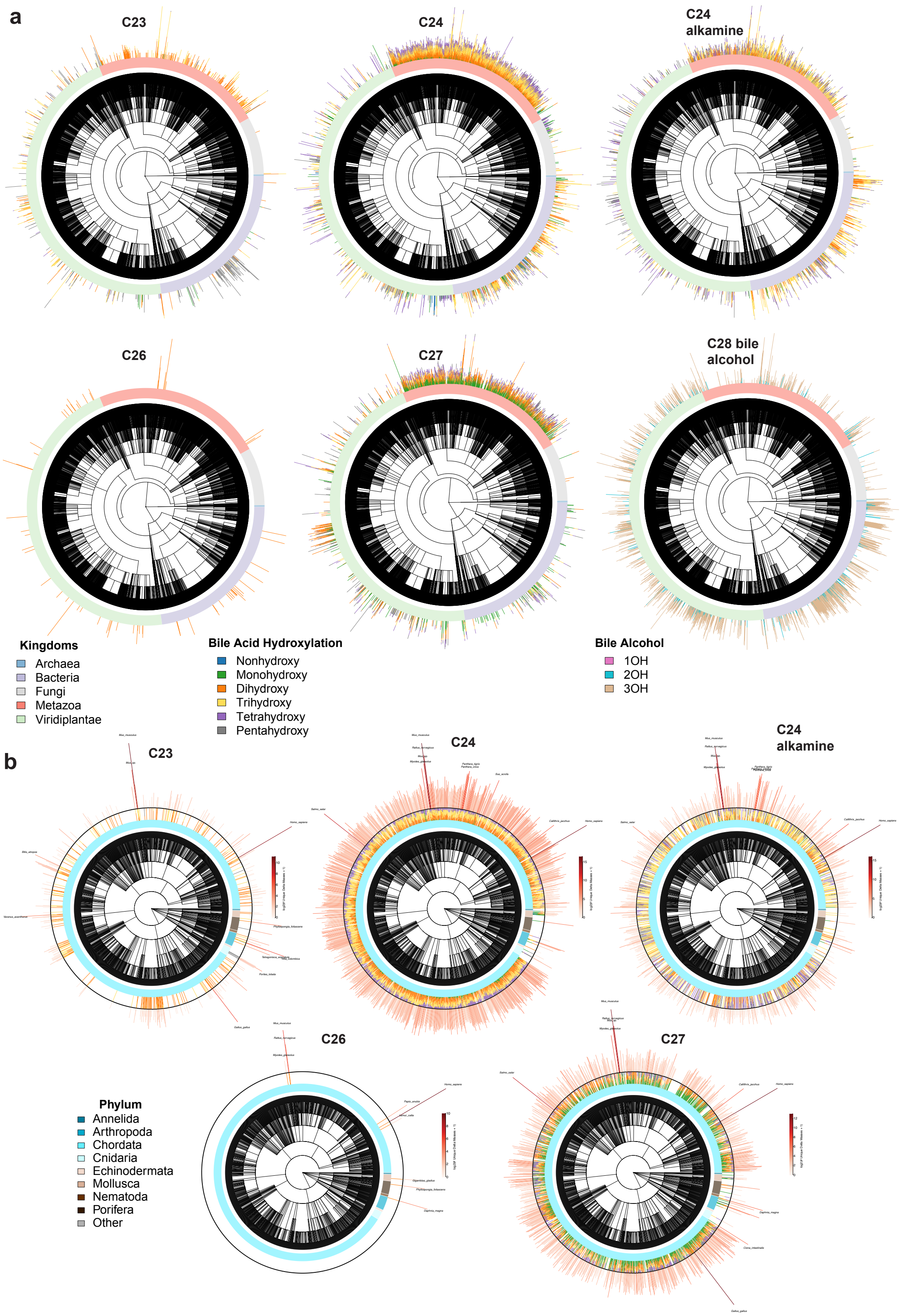


**Supplementary Figure 4. Taxonomic distribution of bile acid core classes and modifications.** Species-level taxonomic trees show the biological distribution of spectra containing the indicated bile acid cores; only taxa with complete taxonomic lineages and entries with positive delta masses were included. Spectra assigned to potential other ion forms are not included. **(a)** Trees spanning all represented kingdoms. The outer annotation ring indicates kingdom, and colored stacked bars show the number of spectra assigned to each hydroxylation subclass. Colors distinguish hydroxylation states for bile acids and the number of hydroxy groups for C28 bile alcohols. **(b)** Metazoa-specific trees. The annotation ring indicates phylum, colored stacked bars show subclass-specific spectrum counts, and the outer red bars represent the total number of unique positive delta masses detected for each taxon, displayed as $\log_2$(number of unique delta masses + 1). Apparent differences in bile acid diversity among taxa may also reflect unequal sampling depth and data availability.

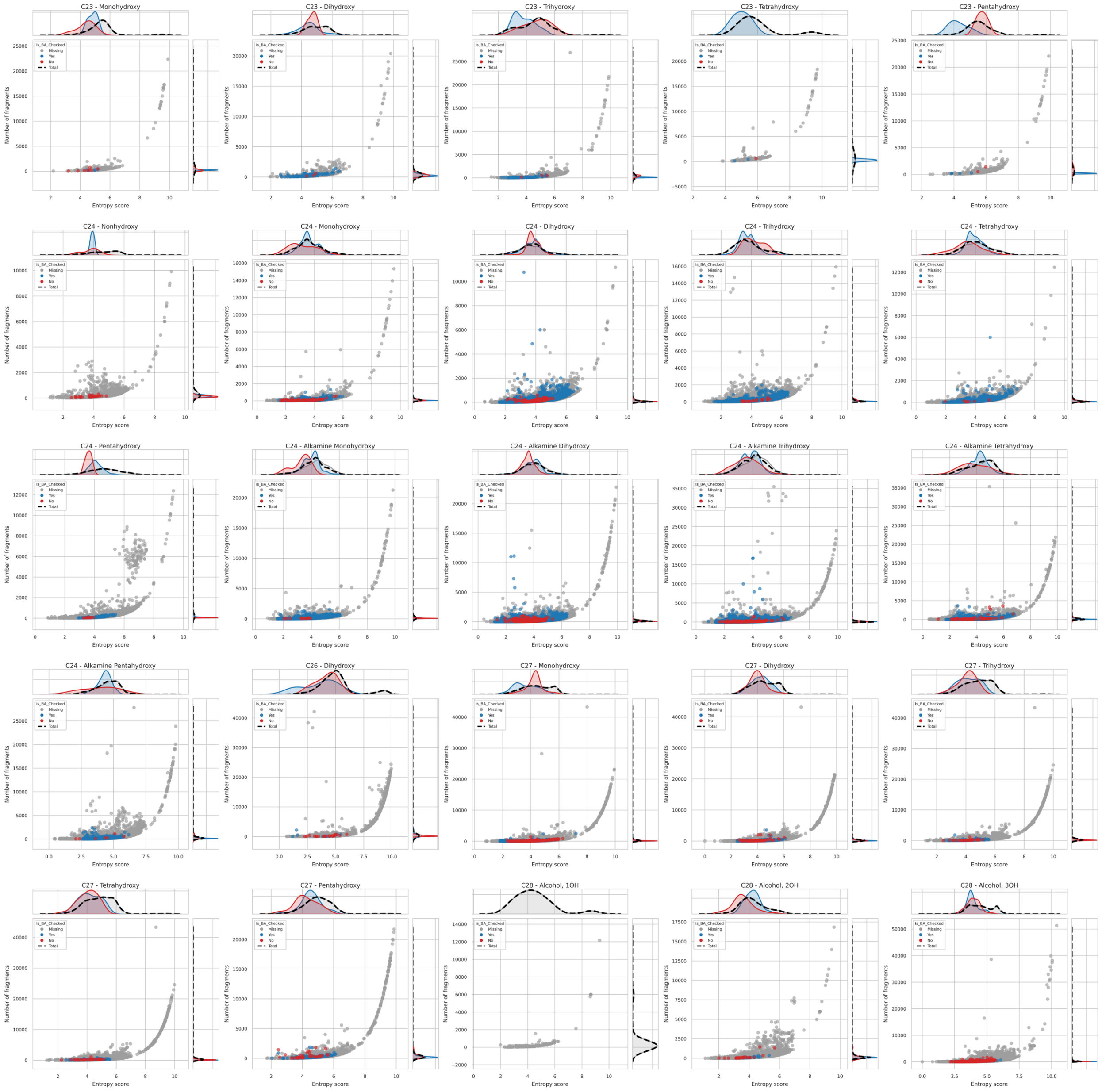

**Supplementary Figure 5. Spectral entropy and fragment-ion counts across bile acid subclasses.** Each panel shows the spectral entropy score and number of fragment ions for spectra assigned to the indicated core and hydroxylation subclass. Scatters are colored according to the bile acid annotation field: red means it had a spectral match to a tetracyclic core structure, blue means it matched to other classes, and gray means it did not have a library match. Marginal density plots show the entropy and fragment-count distributions.

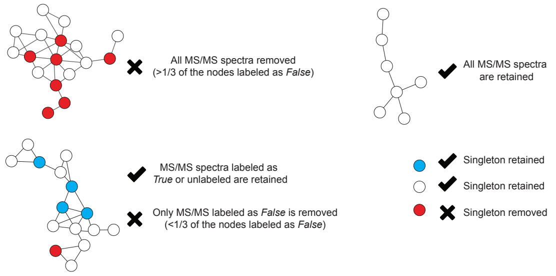


**Supplementary Figure 6. Network-based pruning of molecular families.** Nodes are MS/MS spectra colored by GNPS2 library annotation: blue, bile acid or steroid (True); red, other compound class (False); white, unannotated. Families with more than one-third of annotated nodes labeled False were removed entirely; otherwise, only False nodes were removed. Singletons were retained unless labeled False.

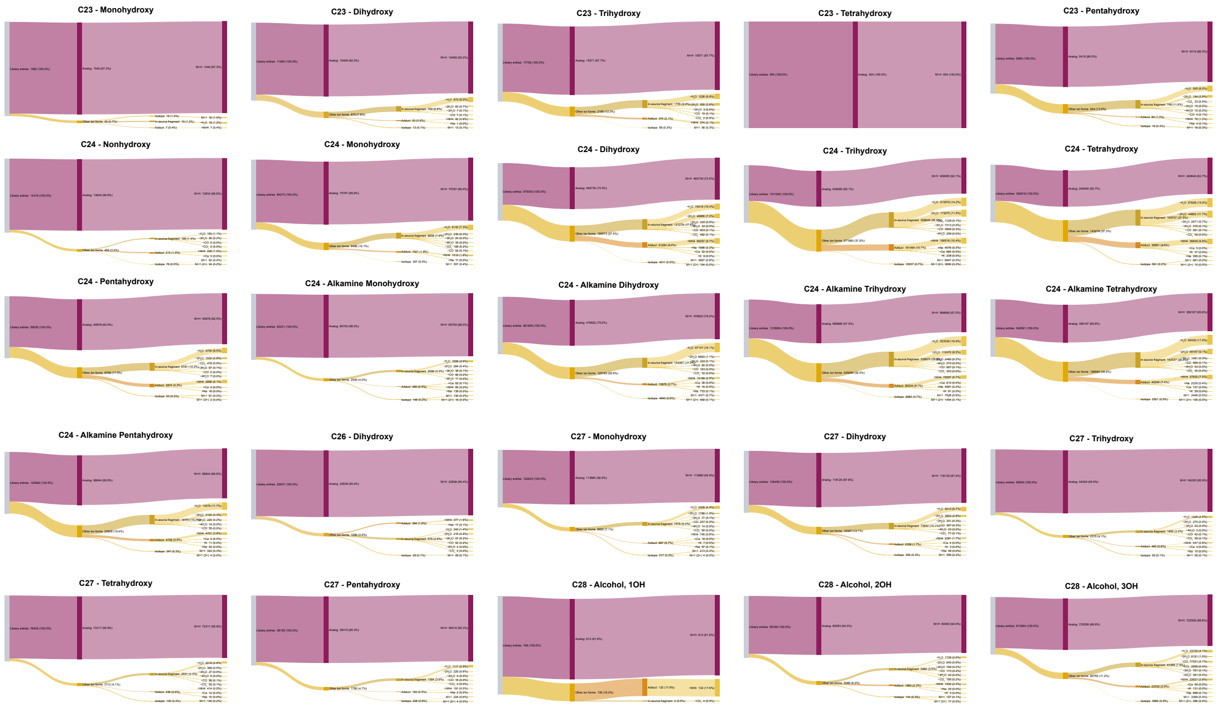


**Supplementary Figure 7. Estimated alternative ion forms within bile acid subclasses.** Sankey diagrams summarize the estimated ion forms for each combination of bile acid core and hydroxylation state. Entries are separated into unflagged protonated analogs and potential alternative ion forms inferred from co-elution and characteristic exact-mass offsets. Alternative forms are further classified as in-source fragments, adducts or isotopologues and assigned to specific forms.

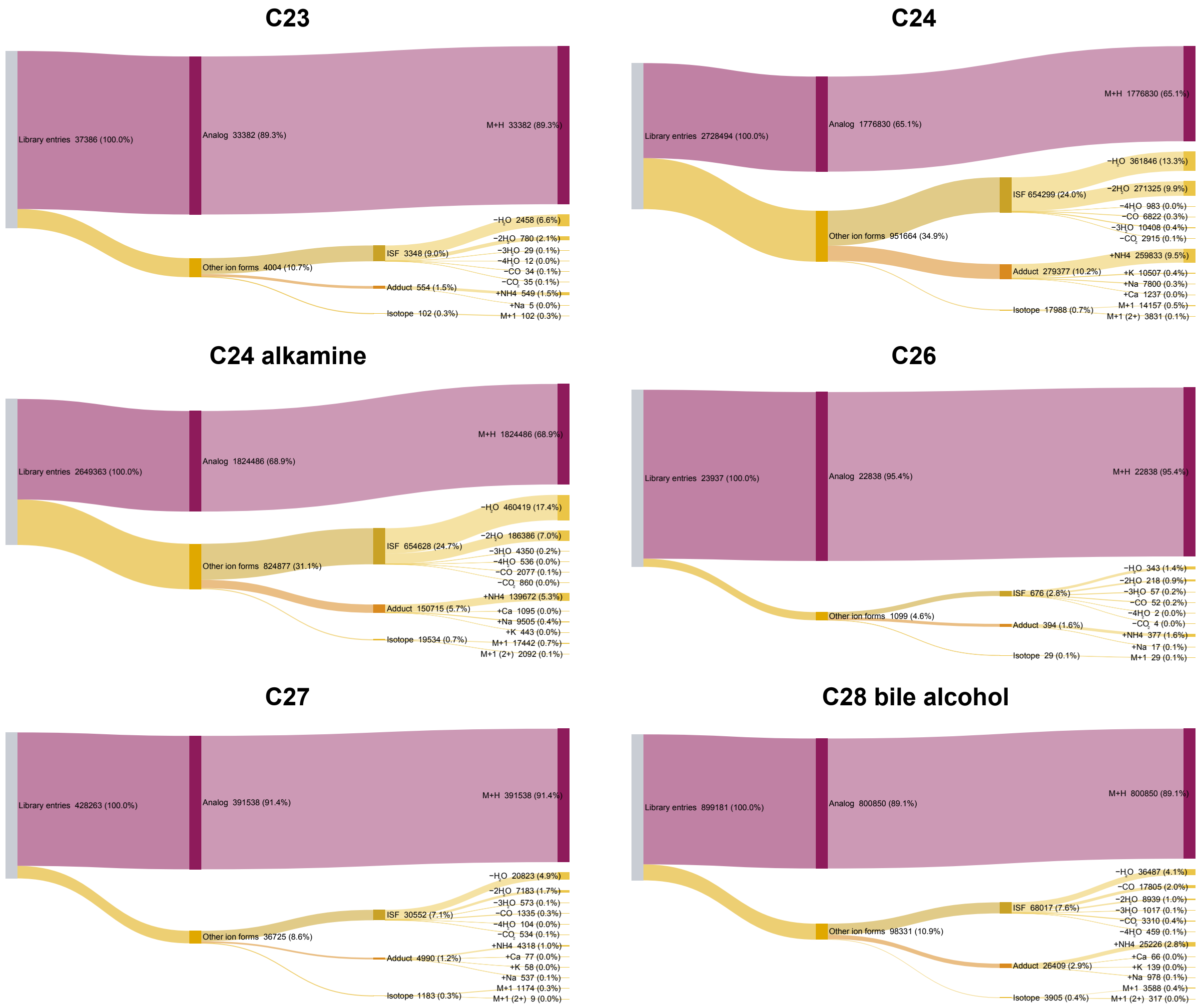


**Supplementary Figure 8. Estimated alternative ion forms across bile acid core libraries.** Sankey diagrams aggregate the estimated ion-form composition of the C23, C24, C24 alkamine, C26, C27, and C28 bile alcohol libraries across hydroxylation states. Entries are separated into unflagged protonated analogs and potential in-source fragments, adducts and isotopologues, with the corresponding ion forms indicated.
